# Monkeys engage in visual simulation to solve complex problems

**DOI:** 10.1101/2024.02.21.581495

**Authors:** Aarit Ahuja, Nadira Yusif Rodriguez, Alekh Karkada Ashok, Thomas Serre, Theresa Desrochers, David Sheinberg

## Abstract

Visual simulation — i.e., using internal reconstructions of the world to experience potential future versions of events that are not currently happening — is among the most sophisticated capacities of the human mind. But is this ability in fact uniquely human? To answer this question, we tested monkeys on a series of experiments involving the ‘Planko’ game, which we have previously used to evoke visual simulation in human participants. We found that monkeys were able to successfully play the game using a simulation strategy, predicting the trajectory of a ball through a field of planks while demonstrating a level of accuracy and behavioral signatures comparable to humans. Computational analyses further revealed that the monkeys’ strategy while playing Planko aligned with a recurrent neural network (RNN) that approached the task using a spontaneously learned simulation strategy. Finally, we carried out awake functional magnetic resonance imaging while monkeys played Planko. We found activity in motion-sensitive regions of the monkey brain during hypothesized simulation periods, even without any perceived visual motion cues. This neural result closely mirrors previous findings from human research, suggesting a shared mechanism of visual simulation across species. In all, these findings challenge traditional views of animal cognition, proposing that nonhuman primates possess a complex cognitive landscape, capable of invoking imaginative and predictive mental experiences to solve complex everyday problems.

---

Consider the following scenario: it’s a Monday morning, and you’re driving to work. You’ve taken this route dozens of times and can navigate without a second thought. Yet today, as you approach your destination, you encounter construction blocking your usual path. You stop, and start to think: “Maybe I could turn right here and head north on a parallel road? Although, that does lead to a one-way street. Perhaps I’ll have better luck if I take a left? That should take me past the bakery, around the school crossing, and eventually direct me back to my destination”. In just a few seconds, you have managed to simulate potential alternate paths and chart a new course for your journey.

This process of problem solving via “mental simulation” takes place entirely in your head, without the need to move a single muscle. You can imagine how things might play out in the future to help you arrive at a solution. Importantly, you need not actually experience the things you are simulating (or their potentially negative consequences, such as going down a one-way street). Your internally generated mental recreations are sufficient to guide your actions. When harnessed effectively, mental simulation is one of the most sophisticated and useful cognitive capacities at your disposal.

Mental simulations can take various forms. For instance, past research has shown that imagining bodily movements relies on a form of mental simulation, as indicated by the overlap in neural circuits involved in imagination and execution of motor actions. This phenomenon is referred to as “action simulation”^1^. Similarly, mental simulation strategies can also explain how we predict outcomes in physical scenes, such as the collapse of a Jenga tower^2,3^. In recent experiments, we have built upon previous intuitive physics studies to provide evidence for a specific type of mental simulation called visual simulation^4,5^. As the name implies, visual simulation involves imagined visual representations corresponding to the objects involved in the mental simulation, similar to how action simulation involves imagined motor representations. In other words, visual simulation incorporates a distinctly visual component into the simulation process. Supporting this idea, we have developed a novel task called Planko (Figure 1) and demonstrated that when people are asked to predict the likely trajectory of a ball falling through an obstacle-filled display, their behaviors and eye movements indicate that they are simulating the ball’s path^5^. Furthermore, we have shown that during these simulations of the ball’s trajectory, motion-sensitive brain regions like the middle temporal area (area MT) respond as if the ball’s motion were being literally seen, even though the stimulus remains static throughout the simulation^4^. Collectively, these findings suggest that humans are indeed capable of visual simulation, and the neural correlates of this process can be observed in visual brain areas.

**Figure 1:**
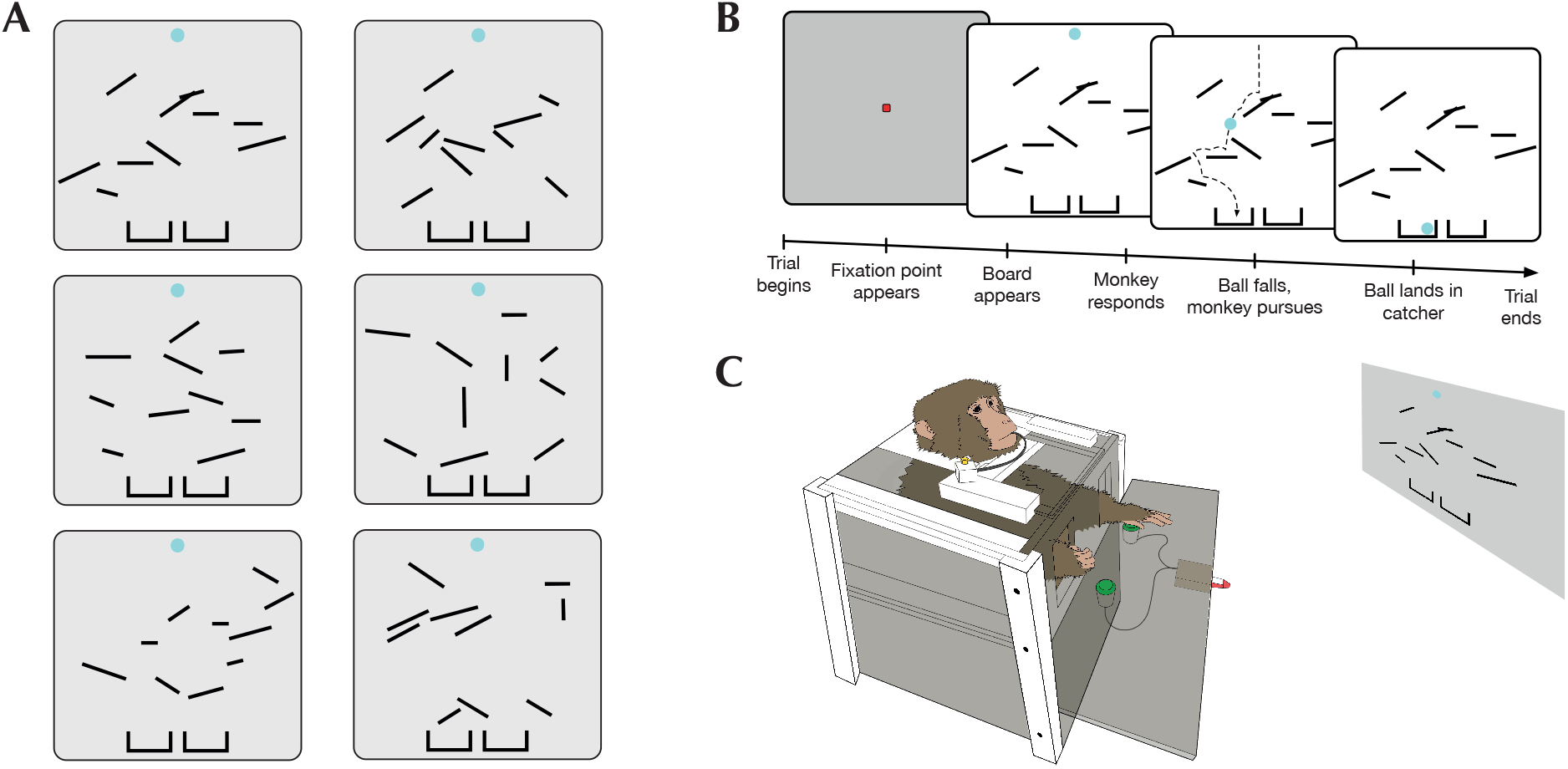
**<u>A</u> -** Examples of Planko boards used in the task. Monkeys were required to predict which catcher the ball would land in, if dropped. In the present example, the three boards in the left column lead to the left catcher, and the three boards in the right column lead to the right catcher. **<u>B</u> -** A schematic of one complete trial, including the pre-response period when the monkeys could potentially simulate the ball’s trajectory and the post-response period when they saw the ball fall. **<u>C</u> -** A diagram of the NHP upright rig setup that was used for training and behavioral testing on the task. Monkeys indicated their responses using one of the two provided buttons, and were given juice reward for correct responses.

Despite the growing body of behavioral and neuroimaging work on simulations in the brain, several questions about the underlying neural mechanisms that support these phenomena remain. Why is this? We suggest that a major obstacle to progress derives from the lack of a compelling animal model. The absence of research on mental and visual simulation in animals is not surprising, given the complexity, introspection, and subjectivity associated with these phenomena. While some recent evidence suggests that computational models of simulation align with nonhuman primate behavior, it remains unclear whether animals are capable of mental simulation, let alone visual simulation^6^. In our current experiments, we aimed to address these questions by replicating our human studies on visual simulation with nonhuman primates (NHPs). We found that when macaques play Planko, their behavioral patterns can be accurately accounted for by models assuming a simulation strategy. Using awake monkey fMRI, we further discovered that when monkeys engage in a simulation of the ball’s trajectory, motion-sensitive brain regions become active, indicating an explicitly visual aspect of the simulation process. Previously, we provided evidence for these same findings with human participants. Together, these results demonstrate that monkeys not only possess the ability for visual simulation but also share the biological foundations of this capability with humans and other nonhuman primates.

## Can monkeys play Planko?

We have developed a task paradigm called Planko to probe visual simulation. During the task, participants are shown displays (also referred to as boards) like the ones shown in Figure 1A and asked to predict which of the two bottom “catchers” the ball will end up in, were it to be dropped through a field of randomly arranged “planks”. While the participants make this prediction (i.e., during the “pre-response period”), the ball remains completely static, suspended in place. Once the participants indicate their choice with a button press, the ball is then in fact dropped during the “post-response period”, thus providing participants with feedback on their responses. While we have been successful in using this task to probe simulation in humans^5^, we wondered if monkeys would be capable of learning it, and if so, would they, too, rely on visual simulation?

We thus set out to train two monkeys (referred to here as Monkey G and Monkey A) to play Planko. We started with extremely simple displays, containing very few planks (Figure 2B). During the early stages of training, we also introduced a “shadow ball” which would reveal a persistent, light gray trace of the ball’s trajectory on certain trials at random (Figure 2A). The shadow ball thus gave away the correct answer on these trials but was nonetheless a useful tool for demonstrating to the monkeys what they were expected to focus on (i.e., the ball’s trajectory). Over time, we gradually increased the number of on screen planks while simultaneously reducing the proportion of trials containing a shadow ball, as well as fading away the shadow ball such that it would reveal progressively lesser amounts of the ball’s trajectory (Figure 2A, 2B). We assessed training success by analyzing monkeys’ task accuracy on non-shadow ball trials.

**Figure 2:**
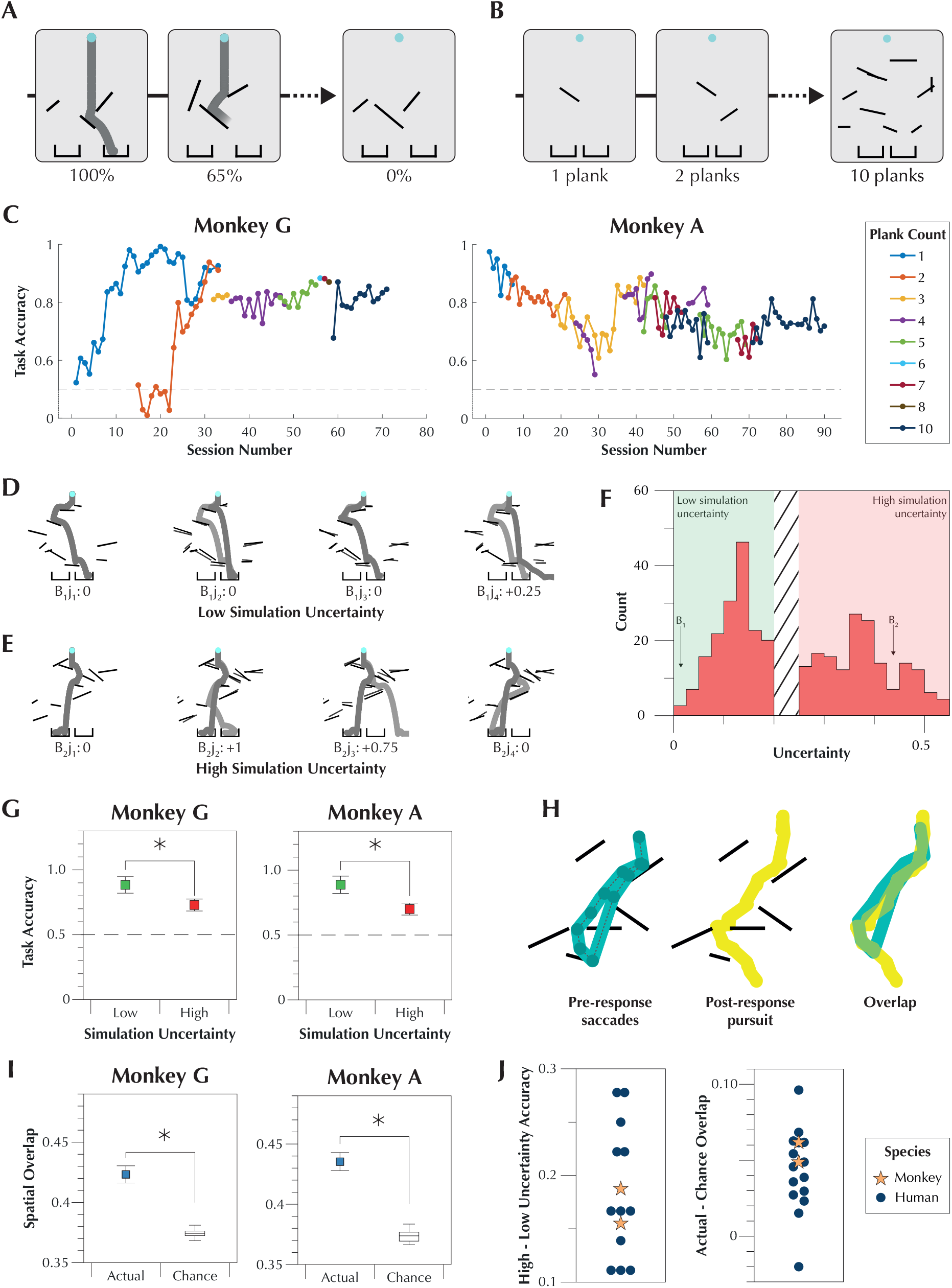
**<u>A</u> -** Examples of the “shadow ball” that was used to train the monkeys on Planko. Shadow ball trials were always intermixed with non-shadow ball trials. Over the course of training, the shadow ball gradually faded away until it revealed none of the ball’s trajectory, in the hopes that monkeys would continue to extrapolate the ball’s trajectory, even when not given. **<u>B</u> -** Examples of various onscreen plank counts that were used during training. Both monkeys began with one plank boards and were gradually progressed along until they were able to navigate ten onscreen planks. **<u>C</u> -** The progression of each monkey’s task accuracy across multiple training sessions in which onscreen planks were progressively increased (non-shadow ball trials only). Both monkeys initially struggled with increasing plank numbers, before arriving at a generalizable strategy that allowed them to maintain consistent task accuracy. **<u>D</u>** - An example of a board where slightly jittering the position of each plank (three jittered examples j_2,3,4_ shown with the original j_1_ underlaid) had a minimal impact on the ball’s final position. Outcome changes were assigned a penalty (in this example, only penalties of 0 were assigned), and used to calculate a simulation uncertainty score. Boards like this one were classified as having a low simulation uncertainty. **<u>E</u>** - An example of a board where slightly jittering the position of each plank (three jittered examples j_2,3,4_ shown with the original j_1_ underlaid) had a significant impact on the ball’s final position. Outcome changes were assigned a penalty between 0 and 1 and used to calculate a simulation uncertainty score. Such boards were classified as having a high simulation uncertainty. **<u>F</u> -** A histogram of all the simulation uncertainty scores assigned to boards from the two monkeys’ task test days. **<u>G</u> -** Task accuracy for Monkey G and Monkey A as a function of simulation uncertainty. Both monkeys were affected by this metric, suggesting that they might be using a simulation strategy. **<u>H</u> -** A schematic depicting the analysis of eye movement overlap between pre-response and post-response trial periods. **<u>I</u> -** Eye movement spatial overlap for Monkey G and Monkey A, relative to a shuffled chance. Both monkeys showed a higher than chance degree of overlap between pre and post response eye movements, consistent with a simulation strategy. **<u>J</u> -** Data from G and I compared to past findings from human subjects. Both monkeys showed behavioral and oculomotor trends that are in line with what we have previously observed in humans (see text for details).

The progression of each monkey’s task accuracy through the aforementioned training (non-shadow trials only) is shown in Figure 2C. Initially, both monkeys struggled with the task whenever we attempted to increase the number of planks on the screen. This is especially apparent in the first 25 sessions. For instance, while Monkey G was able to rapidly learn a one-plank version of the task, his performance fell back down to chance when shown boards with two planks. Similarly, Monkey A demonstrated a progressive reduction in task accuracy as we advanced from one to four onscreen planks. These setbacks suggest that, at least at first, both monkeys relied on strategies that were not robust to changes to the visual properties of the display. This pattern of behavior would not be expected were they relying on simulation to solve the task. Importantly, however, both monkeys eventually modified their approach such that changes to plank number no longer impacted their task accuracy, even when seeing a particular plank count for the very first time. This is most apparent in Figure 2C from session 25 onwards, where we rapidly progressed from two to ten on-screen planks. This newfound invariance to the visual properties of the scene is striking and suggests that the monkeys were able to arrive at a more sophisticated strategy (like simulation) for solving the task (Supplementary Video 1).

## Behavioral Evidence for Simulation in Monkeys

While we were encouraged by the fact that the monkeys were able to play Planko with higher than chance accuracy, that by itself does not provide compelling evidence that they were doing so by using simulation. We thus set out to ascertain whether their behavior on the task was in line with what might be expected, were they engaging in visual simulation. In our previous work, we have shown that human participants’ accuracy on the task depends on the degree of “simulation uncertainty” created by the plank configuration on any board (for details on simulation uncertainty, see **Methods** and Figure 2D/E). In the present study, monkeys were shown boards that fell into one of two discrete simulation uncertainty categories — low, or high (Figure 2F). We predicted that if monkeys were engaging in visual simulation, then their task accuracy would decrease as simulation uncertainty increased. Figure 2G shows both monkeys’ task performance as a function of simulation uncertainty (high vs low). As with humans (Figure 2J), we found that both monkeys were significantly worse at the task when simulation uncertainty increased (Monkey G: t_190_ = 2.75, p < 0.005; Monkey A: t_190_ = 3.27, p < 0.005), suggesting that they were employing a simulation strategy to approach the task.

We also compared monkeys’ eye movements before and after their response on each trial. In the pre-response period, they made saccades while looking at the static image of the board. In the post-response period, they followed the falling ball with smooth pursuit eye movements. Our goal was to determine if the eye movements made while trying to determine the ball’s final position significantly overlapped with the eye movements made while perceiving the ball’s actual falling trajectory (see **Methods** and Figure 2H). We predicted that if monkeys were visually simulating the ball’s movement path to solve the task, then there would be significant overlap between pre-response and post-response eye movements. Figure 2I shows both monkeys’ overlap between pre and post-response eye movements, relative to overlap predicted by chance (see **Methods**). For both monkeys, we found that the degree of observed spatial overlap was greater than chance (Monkey G: t_239_ = 3.49, p < 0.001; Monkey A: t_240_ = 4.17, p < 0.001), suggesting that they may have relied on visual simulation to arrive at the correct answer. This result also precisely mirrors what we have previously observed with human participants (Figure 2J).

## Computational Evidence for Simulation in Monkeys

To develop a computational explanation for playing Planko using simulation and non-simulation strategies, we trained two neural networks: a shallow convolutional neural network (CNN) and a recurrent neural network (RNN) (Figure 3A). We opted for this approach because in previous work, we have shown that such networks (especially RNNs) are capable of spontaneously developing task strategies that resemble visual simulation^7^. The CNN architecture we used in the present study was based on a previous model used in our human research^5^, while the RNN model drew inspiration from neuroscience-inspired motion perception models^8^. Despite inherent architectural differences, we attempted to match the parameters of both networks for consistency. Figure 3B displays the task accuracy of each network when tested on the same boards as Monkey A and Monkey G. Like the monkeys, both networks achieved task accuracy greater than chance (CNN: t_382_ = 15.39, p < 0.001; RNN: t_382_ = 25.27, p < 0.001).

**Figure 3:**
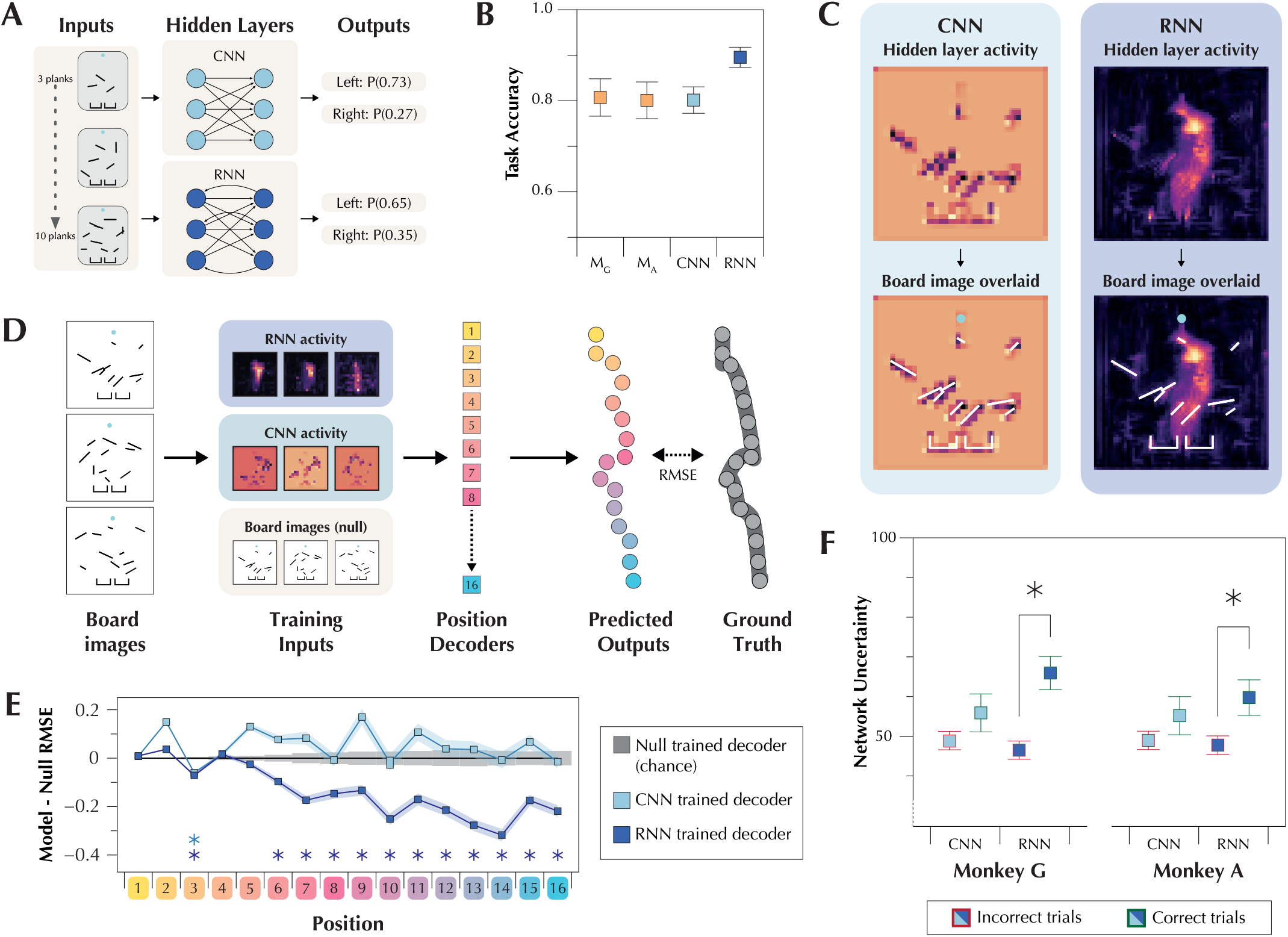
**<u>A</u> -** Examples of two types of networks, a feedforward convolutional neural network (CNN) and a feedback recurrent neural network (RNN) that were trained to solve the Planko task. **<u>B</u> -** Each network’s task accuracy when tested on the same board sets from the monkeys’ task test days. Like the monkeys (M_G_ and M_A_), both networks achieved above chance accuracy. **<u>C</u> -** A heat map showing the average activity of the hidden units on an example board for both the CNN and the RNN. The second row shows the same activity again, but with the input board image overlaid. **<u>D</u> -** A schematic depicting how we quantified whether the ball’s trajectory was represented in the network hidden layer activity. **<u>E</u>** - Average RMSE values for each predicted vs actual position for the CNN and RNN trained decoders, relative to the board image trained (null/chance) model. While the CNN trained decoders almost never achieved greater than chance prediction accuracy, the RNN trained decoders consistently predicted the position of the ball with a high degree of accuracy. **<u>F</u> -** Network uncertainty for the CNN and the RNN as a function of whether each monkey gave the correct or incorrect response on a given board. The CNN’s average network uncertainty was no different for boards that the monkeys got correct vs boards that they got incorrect, whereas the RNN’s average network uncertainty was significantly higher on boards that the monkeys got incorrect compared to boards they got correct.

Having established that neural networks could predict the ball’s final catcher, we aimed to understand the strategy employed by the two networks by probing the activity of their hidden layers (Figure 3C). The top row shows activity maps of both networks, while the bottom row presents the activity maps overlaid with the original board. At first glance, the CNN’s activity predominantly appears to represent the spatial properties of the planks. Conversely, the RNN seems to focus on the ball’s trajectory rather than the planks themselves. Notably, the RNN naturally emerges with this trajectory representation, despite not explicitly requiring or being trained to do so. This observation aligns with the behavior one would expect from a system relying on simulation.

To quantitatively confirm these impressions and investigate whether either the CNN or RNN had represented the ball’s trajectory, we trained 16 position decoders to predict equidistant points along the trajectory. We used the hidden layer activity from each network to assess if these representations contained information about the ball’s path. As a control, we repeated the decoding process using the board images themselves, which explicitly lack path information (Figure 3D). Comparing the decoder outputs to the ground truth trajectory, we found that several of the RNN-trained decoders made better predictions that were closer to the ground truth points than the null-trained decoders. Specifically, significant results were observed for 12 of the 16 decoders after applying Bonferroni correction for multiple comparisons (Figure 3E; see Supplementary Table 1 for detailed statistics). It is also notable that the divergence in prediction ability between the RNN and null-trained decoders appears to be most pronounced towards the latter half of the ball’s trajectory, where the variability in the ball’s hypothetical position is highest.

In contrast, only one of the CNN-trained decoders made better predictions than the null-trained decoders. Instead, most of the CNN-trained decoders made similar predictions to the null-trained decoders (Figure 3E; see Supplementary Table 1 for detailed statistics). These findings confirm our initial suspicion from the activity maps in Figure 3C — that the RNN represents the ball’s trajectory, consistent with a simulation-based approach, while the CNN adopts an alternate strategy based on the statistical regularities of the planks.

Finally, we analyzed whether the boards that caused the highest uncertainty for the networks corresponded to the ones on which the monkeys made mistakes. We did this to determine which network approach, simulation-like (RNN) or not (CNN), best aligned more with monkey behavior. By calculating each network’s board-wise uncertainty (see **Methods**), we found that the RNN displayed significantly higher uncertainty on boards where monkeys responded incorrectly compared to those where they answered correctly (Monkey G: t_190_ = 3.79, p < 0.001; Monkey A: t_190_ = 2.35, p < 0.05). This result suggests that the monkeys struggled with the same boards for which the answer was unclear to the RNN. Conversely, we found that the CNN exhibited no significant difference in certainty between boards that the monkeys got correct compared to the ones they got incorrect (Monkey G: t_190_ = 1.31, p = 0.18; Monkey A: t_190_ = 1.21, p = 0.22), suggesting that the CNN and monkeys struggled with distinct boards. Overall, this result indicates that the RNN’s simulation-like task approach (as depicted in Figures 3C-3E) successfully captures the monkeys’ approach whereas the CNN’s plank analysis approach does not, supporting the idea that the monkeys also engaged in simulation.

## Neural Evidence for Visual Simulation in Monkeys

In the previous experiments, we laid the foundation for the idea that monkeys may possess the ability to engage in simulation. To investigate whether these simulations are indeed visual, we recorded neural responses from the monkeys using an MRI scanner while they played Planko. We achieved this by training the monkeys to play Planko in a setup novel to the monkeys — seated in a horizontal chair in the sphinx position. Our primary hypothesis was that the neural circuits involved in perceiving the falling ball would also be activated during visual simulation. Based on prior human evidence, we predicted that a) motion-sensitive brain regions would be active while the monkeys simulated the motion trajectory of the ball, and b) that this activity would bear voxel-wise pattern similarity to periods when the monkeys actually perceived the ball falling.

To assess these predictions within the MRI environment, we first defined a region of interest (ROI) that was sensitive to motion. We carried out a motion localizer task (see **Methods**) at the beginning and end of each scanning session, in which the monkeys fixated on a central yellow dot while a field of white dots either flickered or moved coherently in the background (Figure 4B). Any brain regions exhibiting stronger responses to moving dots compared to flickering dots, including well-known areas involved in motion processing such as area MT, MST, and V4d (Figure 5A), were deemed motion-sensitive. For subsequent analyses related to the main Planko task, we focused specifically on these motion-sensitive brain regions as our ROI (Figure 5D).

**Figure 4:**
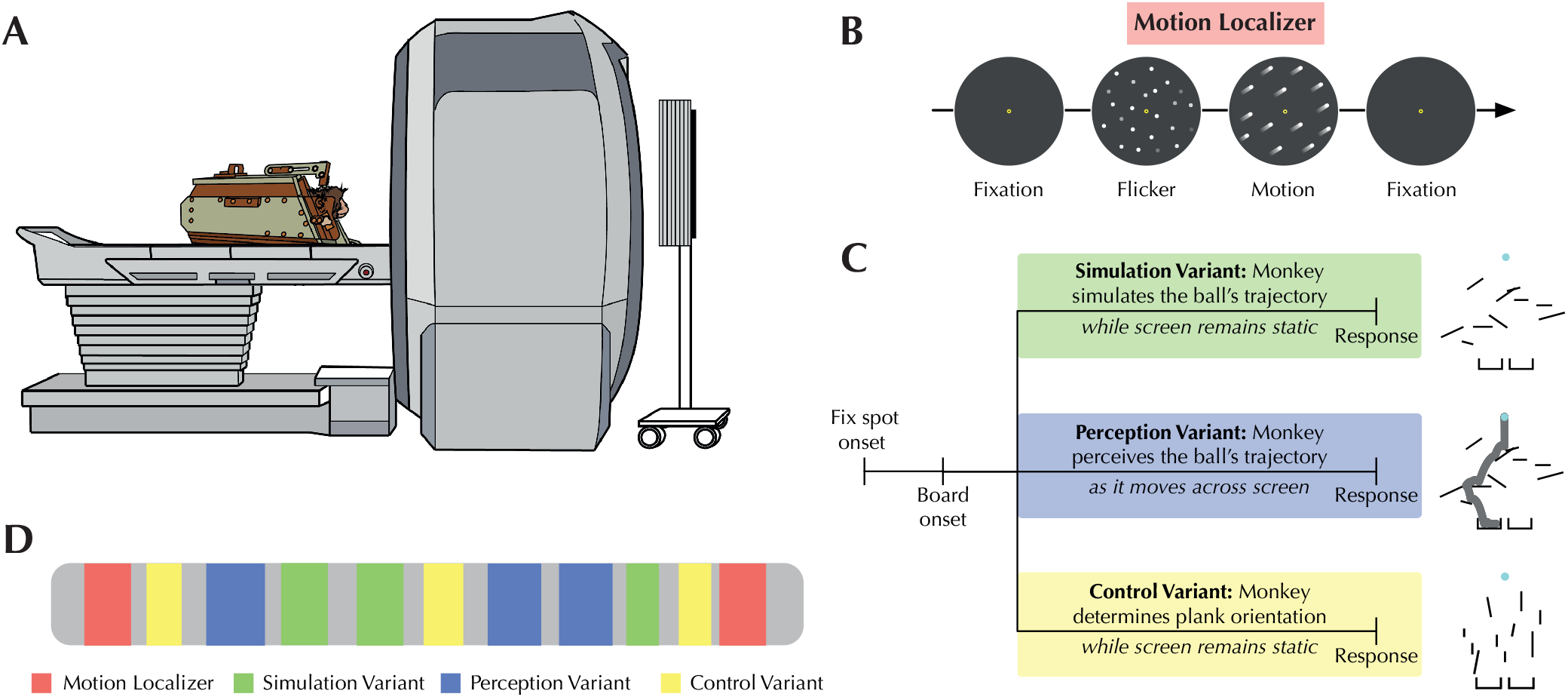
**<u>A</u> -** A diagram of the NHP fMRI setup. Monkeys were seated in the “sphinx” position and placed inside the scanner, where they viewed a screen at the end of the bore and indicated their response using MRI compatible button boxes. **<u>B</u> -** A schematic of the motion localizer task we used to isolate motion sensitive ROIs. **<u>C</u> -** A schematic of the three variants of the main Planko task that monkeys were trained to perform inside the scanner. **<u>D</u> -** An example of one complete scanning session, containing motion localizer blocks at the beginning and end, and several blocks of each task variant randomly interspersed throughout (grey regions indicate interblock intervals).

**Figure 5:**
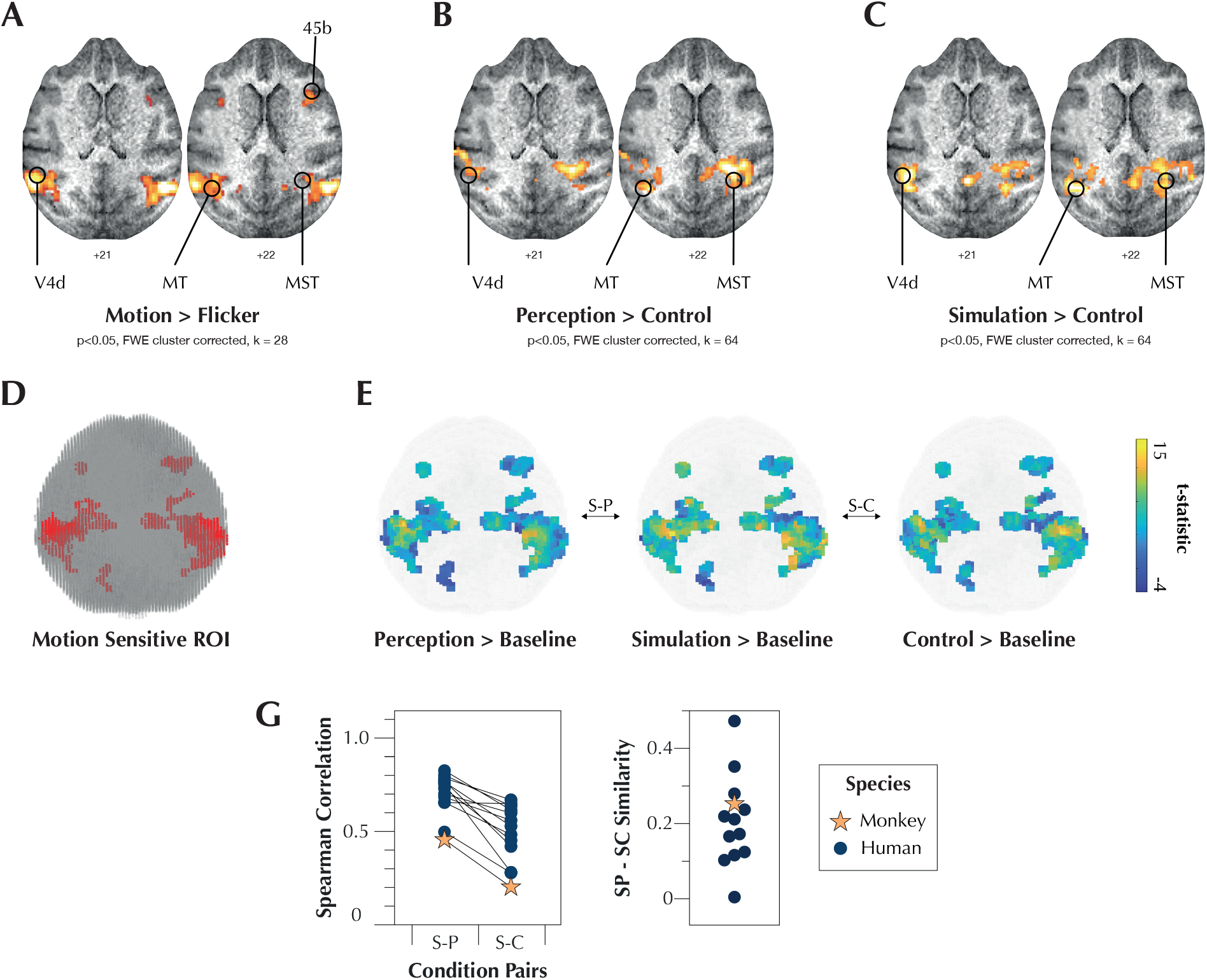
**<u>A</u> -** The result of the Motion > Flicker contrast from the motion localizer task. We observed activity in canonically motion-sensitive brain areas, such as MT, MST, V4d, and 45b. **<u>B</u> -** The result of the Perception > Control contrast from the Planko task variants. Once again, we observed activity in many of the same motion-sensitive areas, such as MT, MST, and V4d. **<u>C</u> -** The result of the Simulation > Control contrast from the Planko task variants. Here too, we observed striking activity in motion-sensitive areas such as MT, MST, and V4d. **<u>D</u> -** A depiction of the motion-sensitive ROI that were used for subsequent representational similarity analyses. All voxels that survived cluster correction at a p < 0.05 FWE threshold were selected. **<u>E</u> -** A schematic showing our main comparisons of interest. We compared the pattern of activity (relative to baseline) in the Simulation condition to both the Perception condition (S-P) and the Control condition (S-C). **<u>G</u> <u>(Left)</u>** - A comparison of S-P and S-C representational similarities. As with the human participants in our previous study (Ahuja et al., 2022), Monkey G’s data also showed an elevated voxel-wise pattern similarity for the S-P comparison relative to the S-C comparison. **(Right)** S-P — S-C similarity for human participants and Monkey G.

To accommodate the spatial and temporal characteristics of the signals acquired in the scanner, we created three distinct task variants of Planko that were administered in a blocked pattern (Figure 4C, 4D). The first variant, referred to as the “simulation variant,” served as the primary experimental condition. In this variant, monkeys were presented with a series of boards and were asked to predict the final catcher for the ball. However, unlike the original Planko task, immediate feedback regarding their choice was not provided. We intentionally removed feedback regarding the ball’s trajectory to ensure that monkeys did not perceive any onscreen motion throughout the simulation block. This design choice was crucial to ensure that any activity related to motion perception would not be mistakenly attributed to simulation. Consequently, we introduced a new “perception variant” that served as a positive control. In the perception variant, monkeys were not required to predict the ball’s trajectory; instead, as soon as the board appeared on the screen, the ball automatically began to fall. The monkeys were then tasked with retrospectively reporting the catcher in which the ball landed. By examining neural responses within our defined ROI from this task variant, we established a template for motion-related activity associated with the ball’s trajectory. This template served as a basis of comparison for the simulation-related activity observed in the previous variant.

Additionally, we developed a “control variant” to serve as a negative control condition. In this variant, monkeys were presented with static boards (similar to the simulation variant). However, unlike the previous variants, monkeys’ responses were no longer dependent on the ball’s trajectory. Instead, the orientations of the planks on the screen were manipulated to be predominantly horizontal or predominantly vertical. Monkeys were required to use the orientation property to provide a response (e.g., pressing left for mostly horizontal or pressing right for mostly vertical). Hence, in this variant, monkeys’ subjective experience during the task closely resembled that of the simulation variant (i.e., making decisions about a static board during naturalistic free viewing). However, the cognitive processes employed were no longer related to predicting the ball’s motion trajectory.

The experimental design described above is consistent with our past human research^4^. Unfortunately, during the course of scanning, one of the monkeys developed a fear response to the MRI machine, which meant that only one of the two animals (Monkey G) adapted to the altered environment of the scanner. The subsequent results thus all belong to Monkey G but are contextualized relative to our previous human findings.

Our results revealed two significant findings. Firstly, within the motion-sensitive ROI, we observed increased activity during the perception variant compared to the control variant (Figure 5B). This finding is expected, since the monkeys viewed the falling ball in the perception variant, whereas no motion was present on screen during the control variant. Strikingly, however, we also found increased neural activity in these same motion-sensitive brain regions during the simulation variant relative to the control variant, despite the absence of any onscreen motion in the simulation variant (Figure 5C). The key distinction between the simulation and control variants lies in the fact that the monkeys engaged in a simulation of the ball’s trajectory exclusively during the simulation variant (and only performed orientation discrimination during the control variant). It thus appears that the act of simulating the ball’s path is capable of eliciting activity in motion-sensitive brain areas, as would be expected in the case of visual simulation. Complete activation coordinates for the motion localizer task and Planko task variants can be found in Supplementary Tables 1 - 3.

Second, we used representational similarity analyses (RSA) to conduct a comparison of voxel-level activity patterns between the simulation variant and both the perception and control variants (Figure 5E). Our goal was to examine whether the activity patterns in the simulation variant exhibited a greater resemblance to those in the perception variant compared to the control variant. Such a pattern resemblance would support the notion that the observed activity during the simulation variant was indeed related to a simulation of the ball’s motion. We found this to be the case, observing a higher pattern resemblance between the simulation and perception variants (r = 0.45, p < 0.001), compared to the pattern resemblance between the simulation and control variants (r = 0.2, p < 0.001) (Figure 5G). Since only one monkey was able to perform the task in the scanner, we assessed the robustness of this result by analyzing how consistent it was with our previous human findings showing evidence of visual simulation. To this end, we pooled the monkey and human results, and then carried out a series of leave-one-out t-tests where one set of observations (either human or monkey) was excluded. Across all iterations, we observed a consistent effect size (d = 1.76; SD = 0.14) irrespective of the identity of the excluded observation, indicating a stable and robust effect that was comparable between humans and monkeys. Additionally, all t-tests yielded significant differences (p < 0.001 for all), further supporting the reliability of our monkey results. Collectively, these findings provide compelling evidence that the simulation of the ball’s trajectory in monkeys evokes perception-like activity in motion-sensitive brain regions, supporting the notion that, like humans, nonhuman primates may be capable of visual simulation.

## Discussion

The results of our study represent significant advancements in understanding nonhuman primate cognition, both behaviorally and neurally. Behaviorally, monkeys demonstrated remarkable proficiency in the complex Planko game, showcasing not just an understanding of its mechanics but also employing a sophisticated strategy beyond mere gestalt-style stimulus response mappings. This behavior also aligns with the performance of a recurrent neural network (RNN) that represents the ball’s trajectory in the activity of its hidden layers, further suggesting that monkeys were engaged in mental simulation. Neurally, we observed notable activation in the motion-sensitive region when monkeys mentally simulated the ball’s trajectory, mirroring the neural activity seen when they saw the ball move – a result consistent with previous human studies. Such similar patterns of neural activity during both perceived and simulated events suggests that monkeys are capable of visual simulation, highlighting the profound capacity of their brains to emulate or predict sensory experiences without external stimuli.

Historically, visual simulation — a cognitive process akin to imagination that can be used to predict and plan for the future — has primarily been studied in humans. Nonetheless, the notion that animals might be capable of *some* form of simulation has gained prominence in recent years. For example, studies on action simulation have suggested that mirror neurons in the motor cortex can internally mimic observed or inferred actions^9,10^, while studies on intuitive physics have demonstrated that when monkeys are asked to intercept a moving virtual ball, their behavior on the task is consistent with a simulation strategy^11,12^. Mazes have also been used to explore simulation in animals. In the visual domain it has been shown that when monkeys are asked to determine the location of the maze’s exit, spatially tuned neurons in parietal area 7a respond with vectors consistent with the path to the solution^13^. Similarly, rodents trained to physically traverse a maze are known to exhibit patterns of neural responses in the hippocampus that reflect the animal’s upcoming path, as if the animals were simulating their future journey^14,15^.

Despite these previous pieces of evidence hinting at simulation abilities in animals, definitive conclusions have been elusive due to the complexity and introspection associated with this cognitive process. Moreover, the simplicity of past paradigms has often left room for alternative interpretations, such as memory recall instead of active simulation. For instance, recent studies revisiting hippocampal replay in rodents question the idea that the observed patterns of neural responses reflect future progression through the maze^16^, especially since such experiments necessarily re-use previously trained mazes. Similarly, in previous maze studies with monkeys, the maze’s exit was generally found within one direction change, making it hard to read out a true, temporally extended simulation. The same can be said about intuitive physics research with primates, as the tasks used in these studies usually involve “simulating” two direction vectors at most, at least one of which is often shown at the beginning of each trial.

In the present study, we addressed several of these limitations. For instance, the configuration of boards on each trial was completely novel, limiting the potential influence of past memory. The ball’s trajectories were also complex, incorporating several motion vectors from start to finish. Finally, no information about the ball’s path was provided (even in the early stages of each trial) leaving it entirely up to the monkey to determine the complete solution. Our findings are thus derived from a more challenging experimental paradigm than used in previous studies and bolster the sparse, existing literature on animal simulation while also introducing the possibility that such simulations are characterized by an imagery-like visual aspect.

The Planko task we have used relies on some understanding of physics, at least at an intuitive level. Based on the current data, we cannot say for sure whether monkeys’ ability to learn the task was driven by prior experience, or an innate core understanding of the physical properties of the real world. Our choice of physics as a testbed, however, was less a matter of emphasizing the role of physics, and more motivated by the need to create a system where simulation could be easily moved between a computational and behavioral space. As a first step, we limited the physical worlds to simple rigid body interactions between a falling ball and a series of static planks. Extending these worlds to include constraints like joints and gears or other dynamic forces could induce even richer internal simulations than those explored here.

Further, if one accepts that humans and other animals carry out internal mental simulations, as our data suggest, then it is important to ask if that holds for simulation environments not rooted in actual physics. Our task could be extended to non-physics based worlds, including arbitrary rule based systems or social interactions. Indeed, from a cognitive science perspective, mental simulation has generally been associated with the ability of one to simulate the thoughts and planned actions of others^17–19^, providing one means by which humans, and possibly other animals, can predict how others might act in a particular situation. While our study focused on predicting the action of an inanimate ball in a highly simplified environment, it remains important to understand if a similar process may underlie monkeys’ ability to predict the thoughts and actions of other animate entities.

Our use of functional neuroimaging in this and a previous study^4^ provide one window into the internal process by which people and monkeys may visually simulate events in the world. This method has many well-known advantages, including its non-invasive nature and its whole brain view. At the same time, it leaves many questions unanswered. Given the spatial and temporal limits of fMRI, we cannot say how detailed the sensory activation in motion areas is. Recordings from populations of the individual neurons in these activated regions could, however, be used to provide a real time readout of the internal state of the simulation in order to directly explore the detailed circuitry supporting this ability.

A significant limitation of the present study was the analysis of neural data from only one animal, despite successfully training two for the Planko task. This issue, while notable, does not substantially detract from the value of our findings, considering the challenges inherent in monkey research and the small sample sizes typically involved. The neural findings observed align closely with past human fMRI research on visual simulation, underscoring the reliability of the results despite the limited data set. Another limitation is the extensive training required for monkeys to achieve high task accuracy. This raises concerns about the impact of experience on the likelihood of engaging in visual simulation. However, training is an indispensable part of teaching cognitive tasks to monkeys. An interesting future research direction could be to explore whether prolonged practice or long-term exposure influence any neural or behavioral effects related to simulation.

In conclusion, our study using the Planko game demonstrates more than monkeys’ understanding of game mechanics; it reveals their capacity for mental simulation to predict outcomes. This insight is a significant leap in comprehending the relationship between the visual brain and mental experience in nonhuman primates. Moving beyond traditional studies focused on simplified tasks, our findings suggest that animal cognition might encompass complex thought processes akin to human experiences, involving contemplation, simulation, or even ‘imagination’ of potential scenarios. This revelation challenges our current understanding of animal intelligence, indicating that monkeys, and possibly other animals, can weave together past experiences, current observations, and future possibilities. It opens new avenues in cognitive neuroscience, hinting at a rich, imaginative mental landscape in the animal kingdom. As we continue to explore these capabilities, we deepen our understanding of the diverse spectrum of intelligence across species, bridging the gap in our comprehension of cognitive processes in the animal world.

## Methods

### Subjects and Surgical Procedures

Two adult male rhesus macaque monkeys (Macaca mulatta; Monkey A and Monkey G) were included in the study. Both monkeys weighed approximately 15kg. Each monkey was surgically implanted with an MRI-safe Peek headpost to help reduce head motion in the behavioral and MRI experimental setups. Surgeries were performed under isoflurane anesthesia, in accordance with the guidelines published in the National Institutes of Health Guide for the Care and Use of Laboratory Animals. Surgical procedures were approved by the Brown University Institutional Animal Care and Use Committee.

### Behavioral Experimental Design

Both monkeys were trained to perform the Planko task (as described in Figure 2A-C). Following training, we carried out behavioral test days with each monkey during which they were shown 192 unique boards over the span of 6-8 blocks from 3 sessions. Each trial began with the presentation of a fixation point, following which a Planko board consisting of one ball, ten pseudo-randomly arranged planks, and two catchers was presented on screen. Monkeys had to determine which of the two catchers, left or right, the ball would fall into, were it to be dropped. Monkeys indicated their choice with a button press. The ball was then in fact dropped, revealing the correct answer. Once the ball landed in its catcher, correct responses were rewarded with a few drops of juice. The proportion of boards on which the ball fell into the left or the right catcher was matched (i.e., 0.565 for each). We used an Eyelink-1000 camera (SR Research) to track monkeys’ eye movements for the entirety of the session. Eye position was sampled at 1 kHz and stored to disk at 200 Hz.

### Simulation Uncertainty Analysis

To explore simulation uncertainty, we modeled the potential for different ball trajectories on each board by introducing positional jitter to the planks and recalculating the ball’s path with a physics engine, as shown in Figures 2D and 2E. Some boards showed significant path deviations with slight plank jitter, while others were unaffected. We used this data to calculate a metric for simulation uncertainty by jittering and recalculating the ball’s path 500 times for each board, then measuring how often the jittered configurations resulted in a different outcome. Boards were then classified as low or high uncertainty based on these outcomes, with the results transformed into a 0-100 scale.

### Eye Movement Analysis

To assess whether monkeys’ eye movements were suggestive of a simulation strategy, we compared their pre-response eye movements (i.e., during hypothesized simulation) to their post-response eye movements (i.e., during perception of the falling ball). It is important to note, however, that eye movements in the pre-response period occurred with a static board presentation, leading to only saccades, while those in the post-response period involved both saccades and smooth pursuit. Due to the distinct nature of saccadic (ballistic) and smooth pursuit (continuous) movements, we did not use traditional oculomotor metrics such as timing and velocity for comparison. Instead, we overlaid the eye movement traces from the pre-response and post-response epochs on top of one another, and then calculated the ratio of the intersection and the union of their areas (Figure H).

Notably, this methodology of measuring similarity does result in some incidental spatial overlap even for eye movement traces that are entirely unrelated to one another. We used this form of incidental spatial overlap to quantify a chance intersection level. We did this by randomly shuffling the post-response eye movements across trials and recalculating spatial overlap on mismatched pairs of traces. We implemented this shuffling protocol for each monkey 50 times and averaged the resulting incidental overlap values on each trial for each iteration. Subsequently, we ended up with a distribution of 50 chance overlap values per monkey. We then compared this distribution to the actual, observed degree of overlap between pre-response and post-response eye movements.

### Deep Neural Network Analyses

We trained a simple feedforward 2-layer convolutional neural network (the “CNN”) and the Index-and-Track (InT) circuit^8^ (the “RNN”) each designed to have around 100K parameters. InT incorporates insights from primate neural circuits implicated in object tracking and has been shown to be more performant and correlated to human behavior compared to vanilla RNNs. The RNN consisted of an input layer, the InT circuit layer and finally the readout layer. The input layer had 64 1x1 convolutional filters, the InT circuit had 64 3x3 recurrent kernels mimicking the lateral connections found in the visual cortex. Finally, the readout was a linear layer that transformed the final RNN hidden state to the classification output. The RNN was trained for T = 24 time steps. The CNN was entirely feedforward with a layer of 3x3 convolutional filters followed by a readout layer similar to the RNN.

Both models were trained using the Binary Cross Entropy (BCE) training objective to classify each Planko board into one of either “left” or “right” classes. Model parameters were optimized with Stochastic Gradient Descent implemented via the Adam algorithm (Kingma & Ba, 2014) with an initial learning rate of 3e-4. Planko boards were of size 64 x 64 pixels with 200K boards for training and 5K boards for testing the models. Training was carried out on a NVIDIA TITAN Xp GPU for 100 epochs while measuring validation accuracy after each epoch over a held-out set of 10K boards.

To test if the models had learnt to represent the ball’s trajectory, we trained 16 position decoders to predict the position of the ball along the trajectory. For both the CNN and RNN, after training the models to classify the boards, their weights were frozen and the hidden state activities elicited by the 200K training boards were recorded. These activities were fed into a model with three layers of 1x1 convolution and pooling operations and finally a linear layer to obtain the final predicted position. For the control (“null trained decoder”), these same decoders were trained to predict the ball positions directly from the 200K Planko training boards. The decoders were trained to minimize the mean squared error between the predicted ball position and the ground truth position derived from the physics engine. Like before, the decoders were trained via Stochastic Gradient Descent and were tested on 5K unseen boards.

Finally, we used the networks’ uncertainty on each board to ascertain which network’s strategy better aligned with monkey behavior. To determine uncertainty, we performed confidence calibration (following training) using temperature scaling^20^. This calibrated probability (P(L) and P(R) for “left” and “right”, respectively) was used to define the uncertainty for a board as 1 - |PL - PR|. Using this measure on both the CNN and the RNN, uncertainty was calculated for each of the boards on which monkey data was collected (the neural networks were not trained on these boards). Finally, the network uncertainty ascribed to the boards was averaged based on whether the monkeys made an accurate response on said board. That is, we asked if the boards with high network uncertainty scores from a particular neural network were also the ones that the monkeys got incorrect, and vice versa.

### Motion Localizer and Planko Task Variant Design

Localizer runs started with a 16-second lead-in period with only a yellow fixation point on screen with a black background. Monkeys fixated on the point for the entire 16 seconds. This was followed by randomly ordered 20-second blocks of white dots that either coherently moved in a given direction (i.e., the Motion condition), or flickered on and off (i.e., the Flicker condition). During the Motion and Flicker conditions, the yellow fixation point remained on screen, and monkeys were required to continue fixating (while ignoring the white dots in the background). The white dots were presented in a circular area with a radius of 6 degrees visual angle around the yellow fixation point. White dots were 0.07 degrees visual angle in size and had a density of 69/degrees^2^. During the Motion condition, the white dots moved at 5 degrees/second, randomly changing direction once per second. Monkeys were rewarded for maintaining fixation, which they did for the entirety of each localizer block.

Task variant runs (Simulation, Perception, and Control) were broken down into blocks of 32 trials each, starting and ending with a 16-second fixation period. Task variant identity was cued by the color of a fixation spot that was presented at the start of each trial. Monkeys were successfully able to task switch between variant types, even within a single session.

### fMRI Scanning Procedures

Monkeys were positioned in an MR-safe chair in the “sphynx” stance, with heads secured using a surgically implanted headpost affixed to the chair’s arm. To minimize movement, the chair was padded with Polyethylene foam. Two floor buttons enabled the monkeys to register their task responses. During the task, monkeys wore earplugs to counteract MRI background noise.

Prior to scanning, monkeys were administered a contrast agent (MION) intravenously to enhance SNR^21–23^. Imaging occurred on a Siemens 3T PRISMA MRI system using a custom six-channel coil. Each session began and concluded with a T1-MPRAGE anatomical image, followed by functional images captured through a specific gradient-echo echo-planar sequence. A 24-inch MRI safe screen displayed the visual stimuli.

### fMRI Data Analyses

Task activity on Planko variants was analyzed using a General Linear Model. The expected BOLD response during the pre-trial period was modeled using a boxcar regressor from stimulus onset to participant response. This model was adjusted for varying reaction times, ensuring the accurate representation of the BOLD signal for each trial ^24–26^. The first two trials in each run and trials with outlier reaction times were treated as nuisance regressors. Similarly, nuisance regressors for trials with outlier reaction times, six motion estimates (translation and rotation), and run identity were also included in the model. After these were integrated with the HRF-convolved task regressors, beta and t-statistic values for the task variants were obtained.

After having derived activity estimates for all variants, we conducted a Representational Similarity Analysis (RSA) to compare variants to one another. In this study, we used voxel-wise t-statistics for each variant (contrasted against baseline) within a motion sensitive ROI as the activity estimates due to their demonstrated reliability for RSA ^27^. We chose the Spearman correlation as our similarity metric, calculating the degree of similarity between the Simulation and Perception conditions (S-P), as well as the Simulation and Control conditions (S-C). The observed S-P and S-C similarities were then directly compared to one another.

## Acknowledgements

We would like to acknowledge Dr. Michael Worden, Dr. Michael Paradiso, Ryan Miller, Kati Conen, and Matthew Maestri for their contributions to this project. We also thank the Brown University Center for Animal Resources and Education for providing animal care, as well as the Magnetic Resonance Imaging Research Facility for their assistance in implementing scanning procedures. This work was supported by the National Eye Institute, National Institute of Mental Health, Office of Integrative Activities, National Science Foundation, and Office of Naval Research (grants R01EY014681, R21EY032713, 2T32EY018080-1, 5T32MH115895-02, 1632738, and N00014-24-1-2026, respectively). Additional support was provided by the Carney Institute for Brain Science, the Center for Vision Research (CVR), and the Center for Computation and Visualization (CCV). We acknowledge the Cloud TPU hardware resources that Google made available via the TensorFlow Research Cloud (TFRC).

## Author Contributions

A.A. and D.L.S designed the behavioral experiments. A.A. performed all behavioral experiments and data analyses. A.A., D.L.S., and T.M.D designed the neuroimaging experiments. A.A. and N.Y.R performed all neuroimaging experiments and data analyses. A.K.A. and T.S. designed all computational experiments. A.K.A. performed all computational experiments and data analyses. A.A. wrote the manuscript with edits and inputs from all authors.

## Data and Code Availability

The data and code pertaining to these results can be obtained from the authors.

## Competing Interests

The authors declare no competing interests.

## Notes

### Competing Interest Statement

The authors have declared no competing interest.

### Summary of Updates

This version of the manuscript has been revised to update acknowledgements, author affiliations, author contributions, data and code availability statements, and competing interest statements.

