## Supplementary Tables 1 - 3 for "Monkeys engage in visual simulation to solve complex problems"

### Supplementary Table 1

**Table 4.1: Activation coordinates for the Motion > Flicker contrast from the motion localizer task**

| ***CHARM Atlas Designation*** | ***Peak x*** | ***Peak y*** | ***Peak z*** | ***Peak t-value*** |
|---|---|---|---|---|
| Visual area 2 (V2) | 23 | -6 | 13 | 6.04 |
| Dorsal visual area 3 (V3d) | 18 | -8 | 20 | 6.22 |
| Dorsal visual area 4 (V4d) | 21 | -7 | 21 | 8.43 |
| Middle temporal area (MT) | 20 | -5 | 21 | 9.69 |
| Medial superior temporal area (MST) | -18 | -1 | 20 | 5.62 |
| Medial intraparietal area (MIP) | 3 | -6 | 24 | 4.48 |
| Caudal dorsal premotor cortex (PMdc) | 11 | 18 | 29 | 4.96 |
| Area 45b | 13 | 23 | 24 | 7.76 |
| Area TEO | -22 | 1 | 16 | 7.90 |

### Supplementary Table 2

**Activation coordinates for the Perception > Control contrast**

| ***CHARM Atlas Designation*** | ***Peak x*** | ***Peak y*** | ***Peak z*** | ***Peak t-value*** |
|---|---|---|---|---|
| Primary visual area (V1) | -26 | -8 | 14 | 3.92 |
| Visual area 2 (V2) | -26 | -6 | 15 | 6.06 |
| Area IPa (IPa) | -18 | 2 | 14 | 4.42 |
| Dorsal visual area 4 (V4d) | -23 | -2 | 21 | 2.77 |
| Middle temporal area (MT) | -16 | -6 | 22 | 4.77 |
| Medial superior temporal area (MST) | 12 | -1 | 22 | 5.14 |
| Temporal parietooccipital associated area (TPO) | 19 | 5 | 14 | 4.62 |
| Area TEO (TEO) | 28 | 1 | 15 | 5.47 |
| Area TEm (TEm) | 23 | 6 | 10 | 4.18 |
| Floor of the superior temporal area (FST) | 17 | 0 | 15 | 4.74 |
| Middle lateral belt region (ML) | -24 | 5 | 21 | 4.69 |
| Anterior lateral belt region (AL) | -25 | 13 | 15 | 5.15 |

### Supplementary Table 3

**Activation coordinates for the Simulation > Control contrast**

| CHARM Atlas Designation | Peak x | Peak y | Peak z | Peak t-value |
|---|---|---|---|---|
| Primary visual area (V1) | -19 | -1 | 21 | 5.20 |
| Visual area 2 (V2) | -26 | -6 | 15 | 4.12 |
| Visual area 3A (V3A) | 10 | -8 | 23 | 3.57 |
| Dorsal visual area 4 (V4d) | 27 | -2 | 17 | 4.37 |
| Middle temporal area (MT) | -18 | -6 | 22 | 4.96 |
| Medial superior temporal area (MST) | -19 | -1 | 21 | 5.20 |
| Temporal parietooccipital associated area (TPO) | -20 | -2 | 21 | 5.18 |
| Area V23 | 1 | -5 | 20 | 5.58 |
